## Additional data file 1 for "Most human DNA replication initiation is dispersed throughout the genome with only a minority within previously identified initiation zones"

**Fig. S1:** ***Concentration-dependent detection of BrdU in human genomic DNA by nanopore sequencing – further figures, accompanies figure 1.***

**A)** Frequency distributions of BrdU probabilities at every thymidine position for nanopore sequencing reads from hTERT-RPE1 cells grown in the indicated range of BrdU concentrations, for the indicated times. Inserts show the full y-axis for the 50 μM sample. **B)** Frequency distributions of the fraction BrdU incorporated in each window (290 thymidines) for the same reads from hTERT-RPE1 cell cultures as in Supplementary Fig. 1A. Insert shows the full y-axis for the 50 μM sample. **C)** Top three rows: BrdU detection as determined by Mass Spectrometry, DNAscent or BrdU-IP for hTERT-RPE1 or HeLa-S3 cells treated with the indicated time and range of BrdU concentrations (as for Supplementary Fig. 1A, B and HeLa-S3 in Fig. 1). Bottom three rows: Pairwise comparisons of fraction BrdU incorporated in DNA as determined from Mass Spectrometry, DNAscent or BrdU-IP for hTERT-RPE1 and HeLa-S3 cell cultures. Dashed line is y=x. (HeLa-S3 mass spectrometry vs DNAscent shown in Fig. 1E). **D)** Meta-analysis of fraction BrdU detected by DNAscent at thymidine positions relative to CGI centres for reads from HeLa-S3 cells treated with 0.0, 0.3, 1.5, 10 or 50 μM BrdU (as per Fig. 1B, D; 0.0 μM and 50 μM also shown in Fig. 1G). Fraction BrdU calculated in 100 bp windows. Dashed lines (200 bps apart) indicate the minimum bounds of CGIs. CGIs were separated into three groups based on mean methylation level, blue lines show the top third (high methylation), black lines show bottom third (low methylation).

**Supplementary Figure S2:** ***Single molecule detection of DNA replication dynamics on ultra-long nanopore sequencing reads – further figures, accompanies figure 2***

**A)** Idiogram [81] of the human genome (assembly GRCh38) illustrating the sequence coverage of BrdU containing reads (heatmap) and the location of identified replication initiation sites (circles) in the whole genome dataset from HeLa-S3 cells. **B)** Whole genome dataset from hTert-RPE1 cells represented as in panel (A). **C)** Histogram of HeLa-S3 DNAscent initiation site lengths (kbp). Dashed line indicates 5 kbp cut off for high resolution initiation sites (<5 kbp). **D)** Histogram of relative distance between each DNAscent initiation site and the nearest Ok-seq initiation zone (red line) with randomised data (blue line) and confidence intervals shown (dashed grey lines, inner 95%, outer 99%).

**Supplementary Figure S3:** ***DNAscent reveals stochastic replication initiation sites not identified by population studies – further figures, accompanies figure 3.***

**A)** Summary plots and heatmaps comparing replication initiation sites identified by DNAscent, and sorted by replication timing (blue), with: proportion GC content, Ok-seq initiation zones (HeLa-S3) [16], PU-seq initiation zones [17], optical mapping (ORM) initiation zones (HeLa-S3) [37], ini-seq initiation sites [23], ini-seq (updated method) initiation sites and SNS-seq initiation sites from the same publication [24], ORC ChIP peaks (HeLa-S3) [26], Mcm7 ChIP peaks (HeLa) [27], SNS-seq initiation sites (HeLa-S3) [46], G4 K minus, G4 K plus, G4 PDS minus, G4 PDS plus (all yellow; HEK-293T) [82] from HeLa-S3 datasets where available as indicated. Ok-seq, Pu-seq and ORM are also shows in Fig. 2. **B)** Summary plots and heatmaps as in Supplementary Fig. S3A except DNAscent initiation sites were divided into two groups, overlapping with Ok-seq initiation sites (focused, dark blue) or not overlapping (dispersed, green). **C)** Summary plots show the mean signal across DNAscent initiation sites for the heatmaps shown in

Supplementary Fig. S3A, here including comparison to random expectations (black line) and 99% confidence intervals (grey band) from 1000 randomisations of DNAscent initiation site co-ordinates. **D)** Summary plots show the mean signal across focused (blue) and dispersed (green) DNAscent initiation sites for the heatmaps shown in Supplementary Fig. S3B, here including comparison to random expectations (black line) and 99% confidence intervals (grey band) from 1000 randomisations of DNAscent initiation site co-ordinates.

**Supplementary Figure S4:** ***High coverage single molecules at MYC, AFF2 and HBB previously published origins – further figures, accompanies Figure 4***

**A)** Top left: Replication fork direction from published Okazaki fragment sequencing [16] across a 1 Mb region including the MYC gene. The location of Ok-seq determined initiation (dark blue) and termination zones (light blue) are marked. Open box indicates the 54 kb locus targeted for enrichment via nCATS. Left middle: Stacked single molecule DNAscent DNA replication profiles with fraction BrdU incorporation indicated via the heatmap (300 bp windows). Gene annotations and the location of ORM and Ok-seq IZs are marked above the heatmap. Right middle: For each single molecule DNAscent DNA replication profile, computationally determined replication kinetics are shown: forks with direction (arrows), initiation (light green bars) and termination sites (dark green bars). Bottom: Two example single-molecule nanopore sequence reads are plotted with DNAscent BrdU probabilities (black dots) and fractions BrdU incorporated (blue lines) as in Fig. 2. For each read the respective heatmap and replication kinetics are reproduced from stacked profiles above. Top right: Ensemble of DNAscent single molecule replication fork direction across the target enrichment locus. The grey ribbon indicates the 95% confidence limits from 1000 randomisations of individual replication fork directions. **B)** and **C)** show replication fork direction from published Okazaki fragment sequencing, stacked single molecule reads with BrdU incorporation and determined replication kinetics as for **A)** for HBB **(B)** and AFF2 **(C)** loci.

**Supplementary Figure S5:** ***A fifth of initiation sites are focused by high levels of proximal transcription to define initiation zones – further figures, accompanies Figure 5.***

**A)** Replication fork direction relative to the level, direction and length of transcription for HeLa-S3 cells. Top panel for low expressed genes (TPM < 0.408), bottom panel for high expressed genes (TPM > 0.408). Panels from left to right are for quartiles of transcript length from short to long. Individual panels are as presented as in Figure 5A. **B)** Replication fork direction relative to the level and direction of transcription for hTERT-RPE1 cells. The bottom half (left) or top half (right) of transcribed genes were determined by normalised transcript counts (Transcripts Per Million; TPM). Forks identified by DNAscent occurring within 10 kb of transcription start sites are included. The count of replication forks co-directional with transcription is shown in green and counter-directional in blue. Confidence intervals from 1000 randomisations are shown in grey (99%) around average fork density (in black). **C)** As B except forks identified by DNAscent from HeLa-S3 reads occurring within 10 kb of transcription end sites are included. **D)** As B except forks identified by DNAscent from hTERT-RPE1 reads occurring within 10 kb of transcription end sites are included. **E)** Heatmaps comparing replication initiation sites identified by DNAscent with: transcription, DNase-seq, and ChIP-seq for the following histone marks: H2AFZ H3K4me1, H3K4me2, H3K4me3, H3K9ac, H3K9me3, H3K27ac, H3K27me3, H3K36me3, H3K79me2, H4K20me1. Initiation sites are separated by S-phase replication timing (early or late) and whether they overlap with Ok-seq IZs (focused/dispersed). Comparison with focused early sites shown in dark blue, and with dispersed early sites shown in mid blue. Comparison with focused late sites shown in red, and with dispersed late sites shown in orange.

**Supplementary Table S1: BrdU-IP data**

Relative number of reads (uniquely mapping reads and after excluding duplicates) between pooled samples giving semi-quantitative BrdU-IP data of BrdU incorporation for HeLa-S3 and hTERT-RPE1 samples as in Fig. 1 and Supplementary Fig. S1.

**Supplementary Table S2: Fraction nascent reads**

Fraction of reads classed as replicated from HeLa-S3 20hr cultures and hTERT-RPE1 2hr, 24hr, 27hr cultures as shown in Fig. 1 and Supplementary Fig. S1. Reads were counted as replicated where >5% of thymidine positions had a probability ≥0.5.

**Supplementary Table S3: Experimental metrics for single molecule detection of DNA replication dynamics**

**A)** Sequencing metrics for the whole genome sequencing datasets from HeLa-S3 and hTert-RPE1 cells. **B)** The number of leftward and rightward forks, initiation and termination sites identified in the HeLa-S3 and hTERT-RPE1 whole-genome datasets, from all reads or when analysing just those reads where at least one 3000 thymidine window contained >5% thymidine positions with a BrdU probability ≥0.5 (nascent reads). **C)** Sequence and location of guide RNAs used in the targeted enrichment experiments (nCATS). **D)** Sequencing metrics for the nCATS targeted enrichment sequencing dataset from HeLa-S3 cells.

**Supplementary Table S4: DNAscent replication initiation site density (RIGR) for various genomic regions.**

Density (RIGR) of DNAscent replication initiation sites (from HeLa-S3 and hTERT-RPE1 datasets) in various genomic regions.

### Supplemental Figure S1

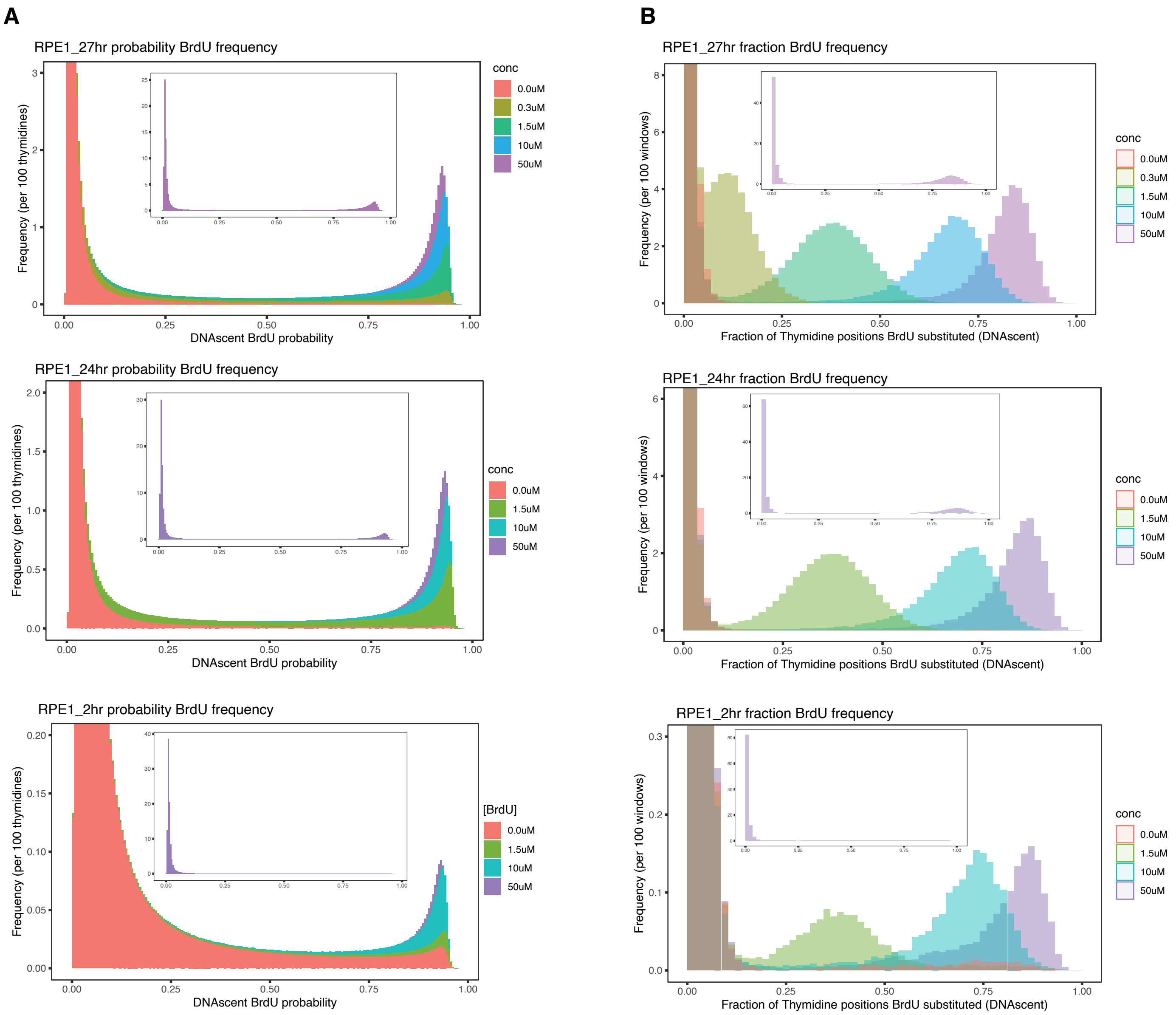

### Supplemental Figure S1

C

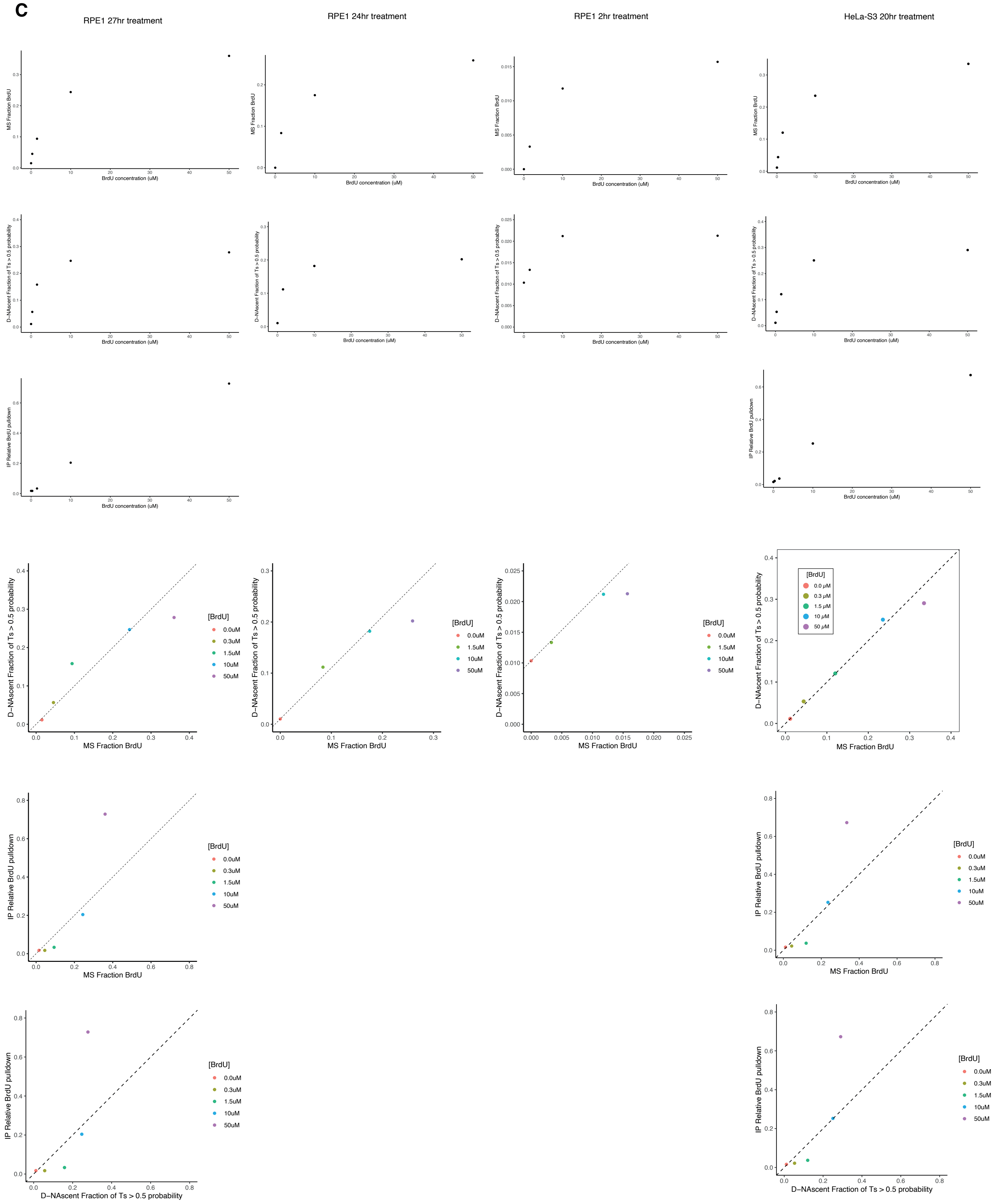

#### Supplemental Figure S1

D

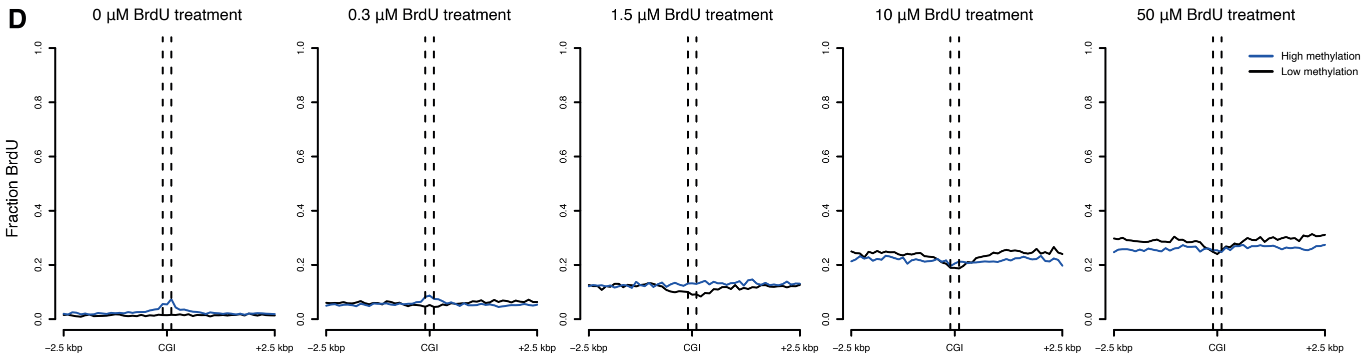

### Supplemental Figure S2

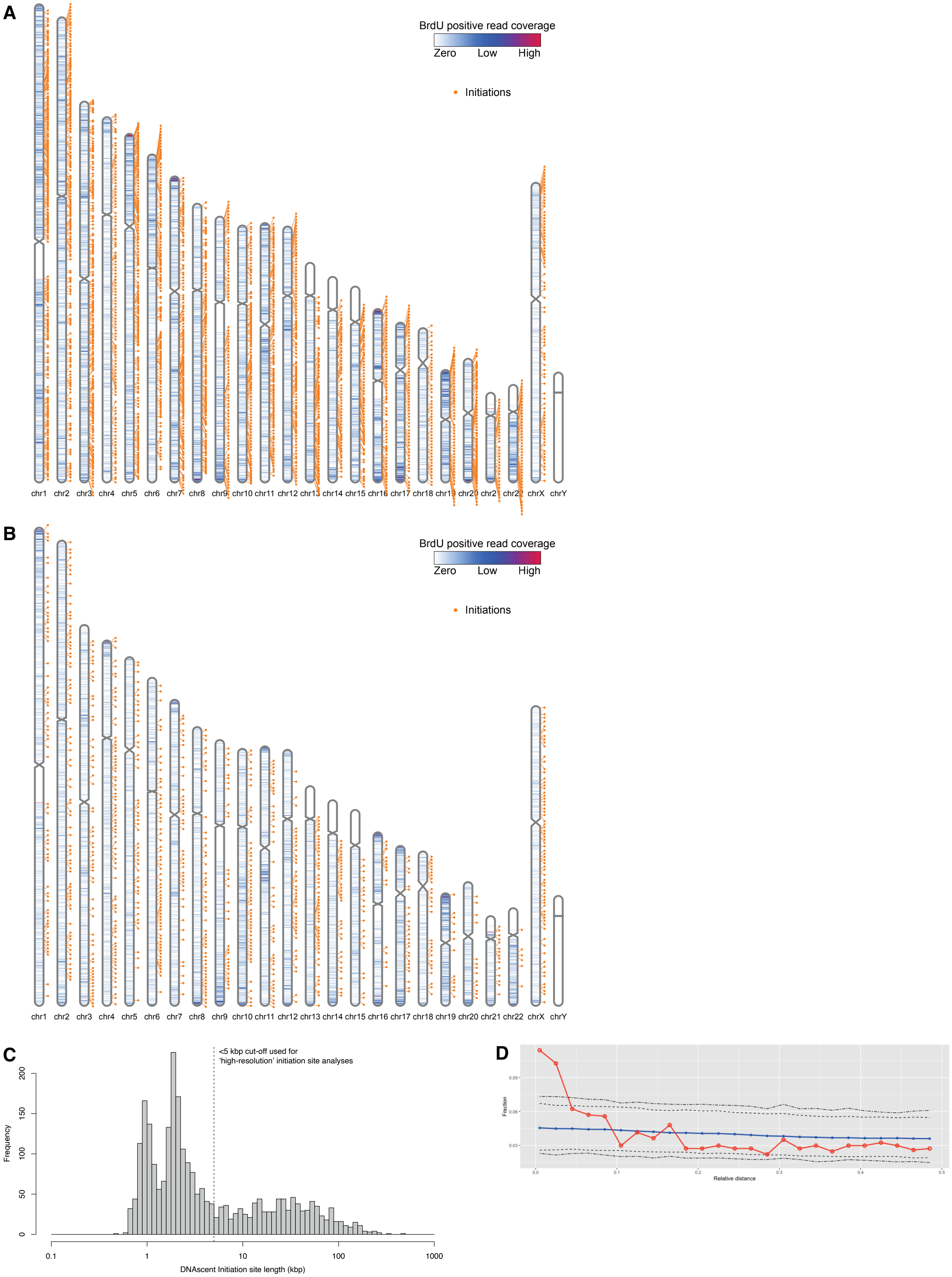

### Supplementary Figure S3

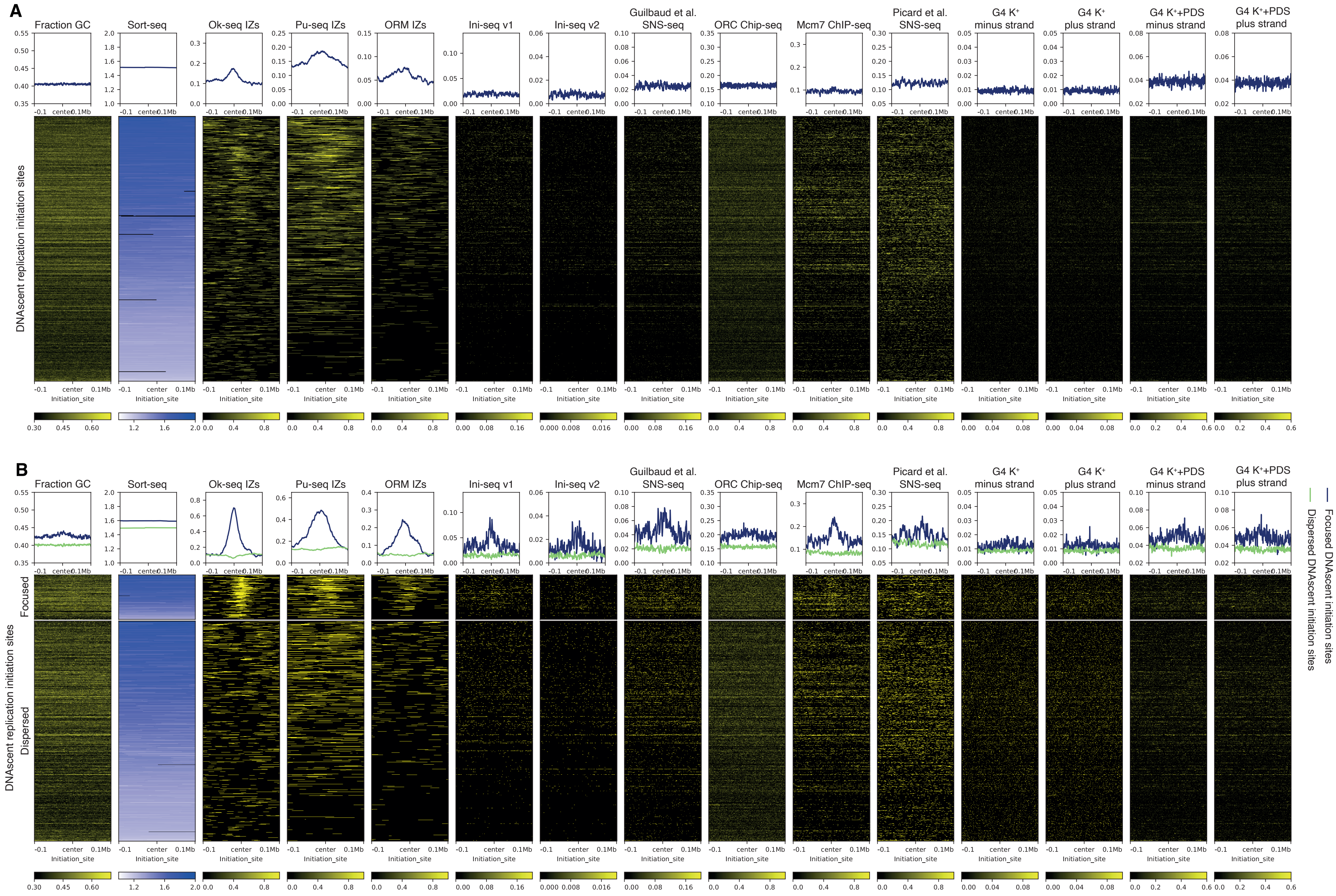

### Supplementary Figure S3

**C**

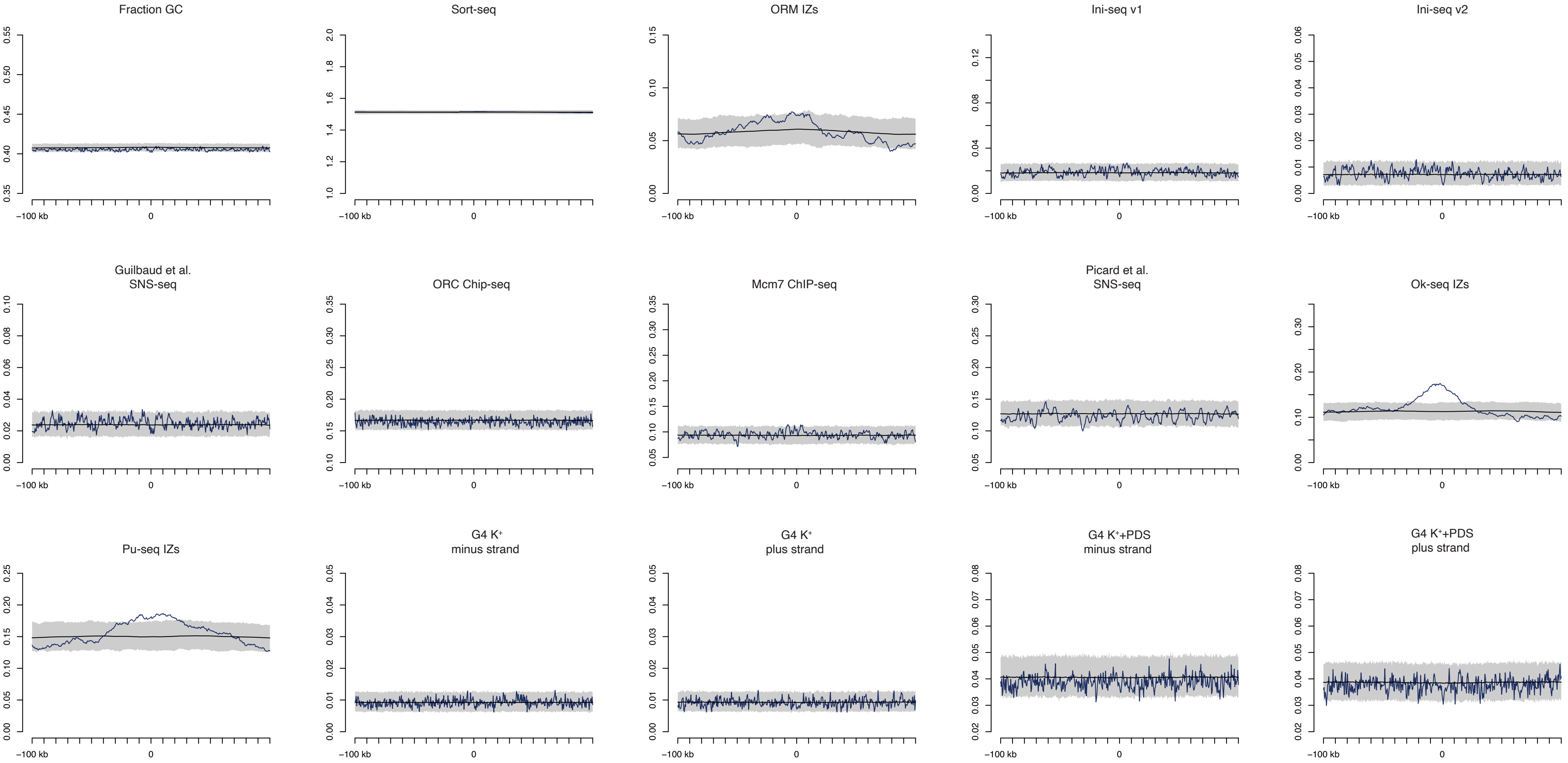

#### Supplementary Figure S3

**D**

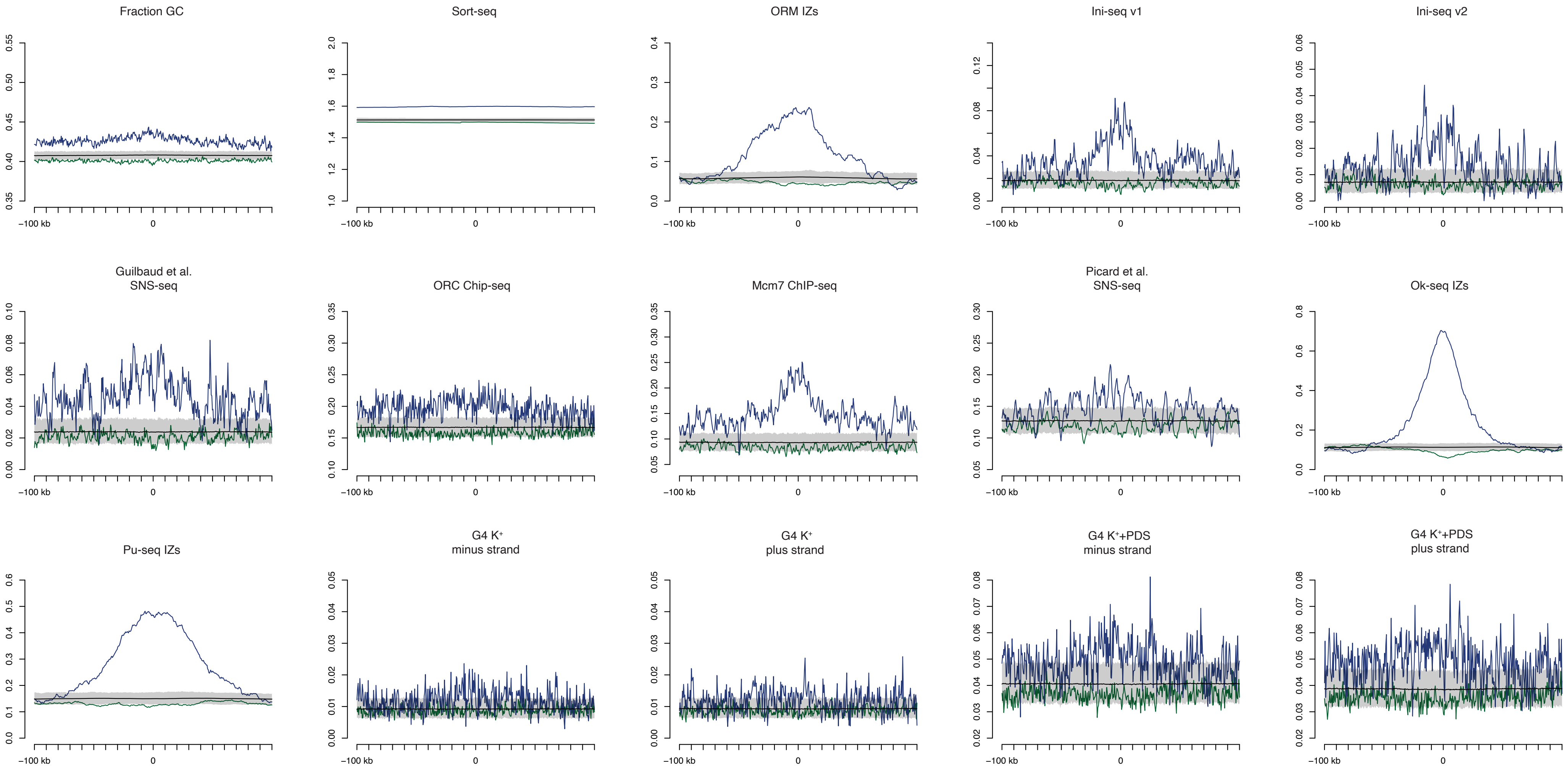

### Supplementary Figure S4

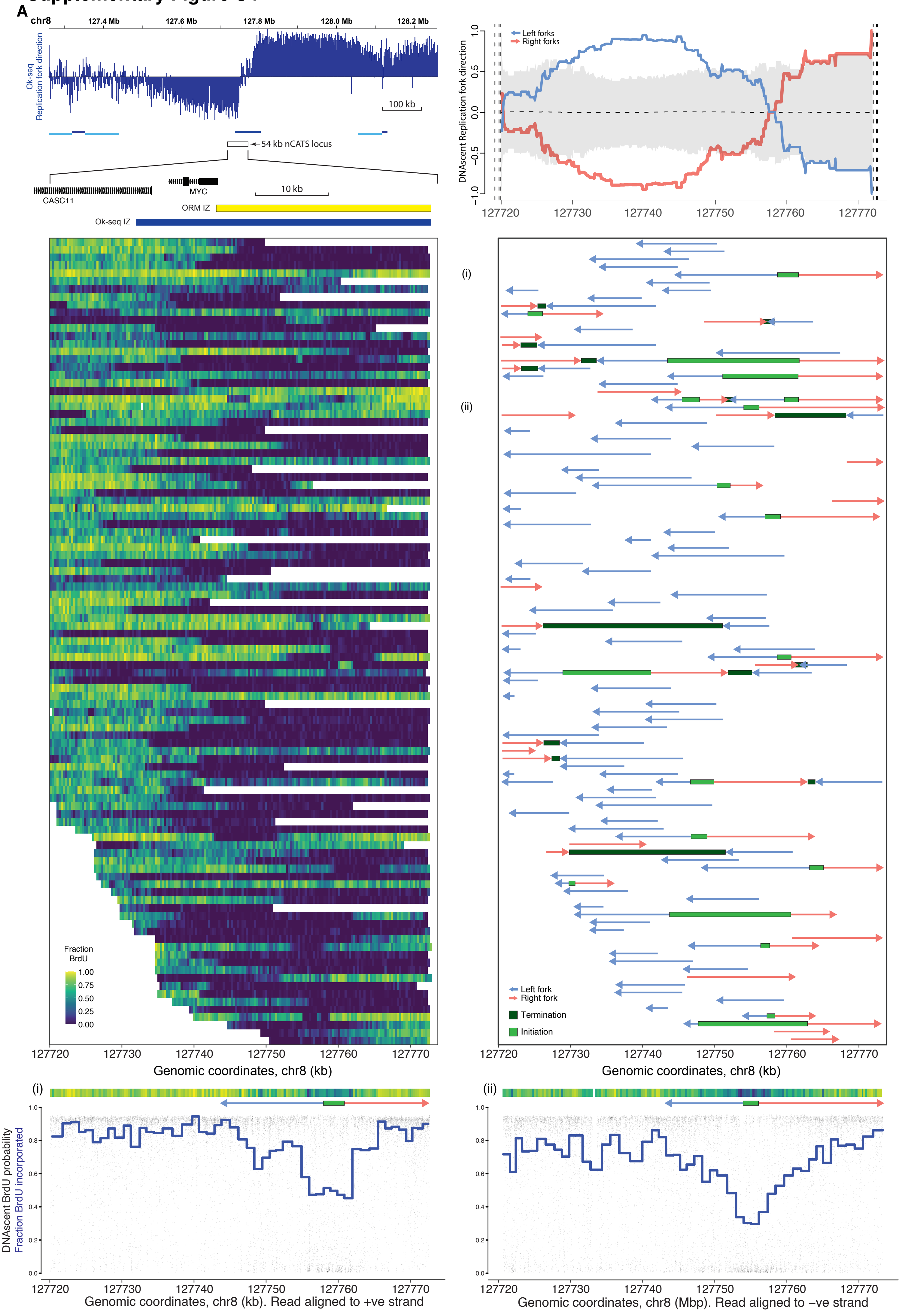

B

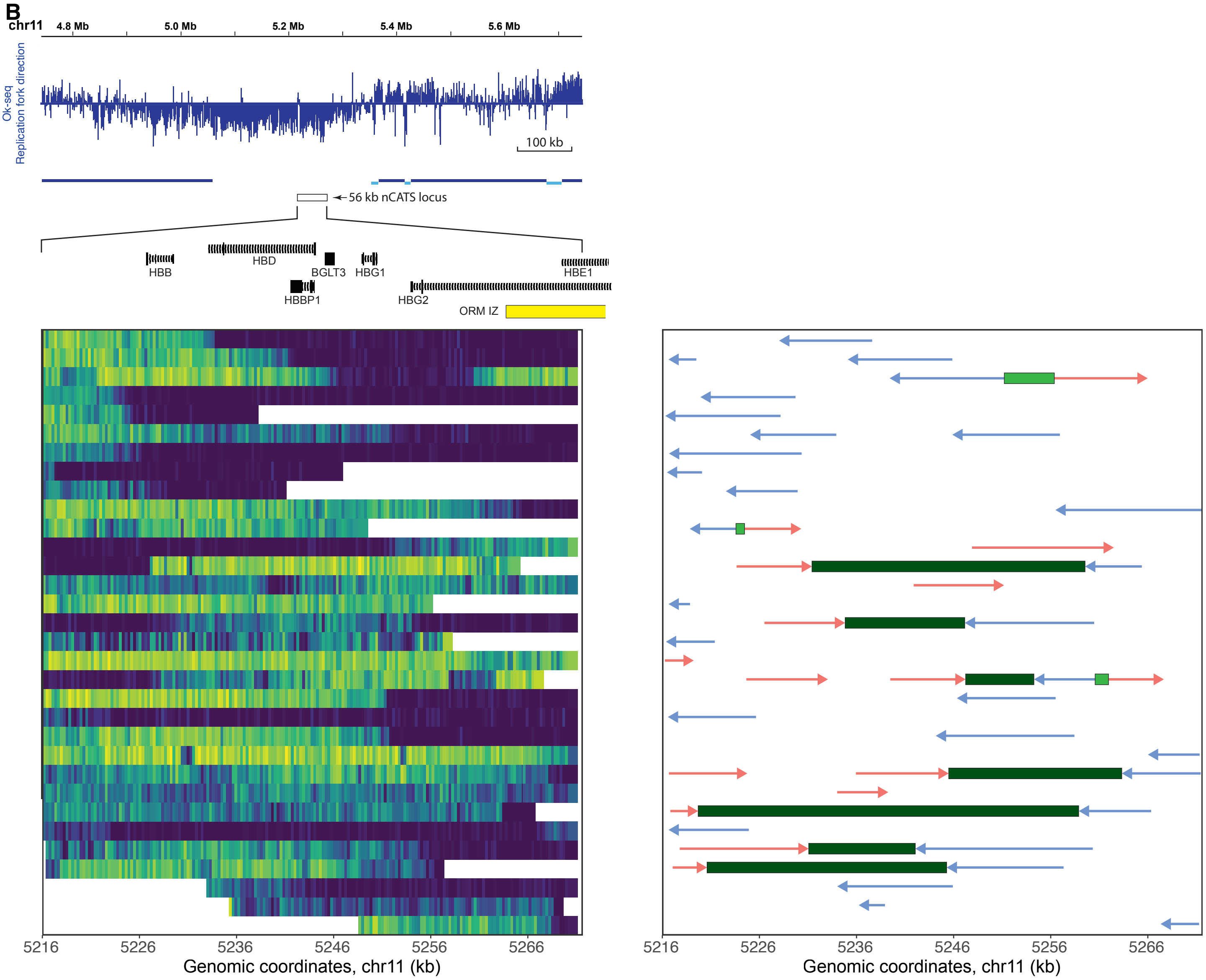

chr11
4.8 Mb
5.0 Mb
5.2 Mb
5.4 Mb
5.6 Mb
Ok-seq
Replication fork direction
100 kb
56 kb nCATS locus
HBB
HBD
BGLT3
HBG1
HBE1
HBBP1
HBG2
ORM IZ
5216
5226
5236
5246
5256
5266
Genomic coordinates, chr11 (kb)
5216
5226
5236
5246
5256
5266
Genomic coordinates, chr11 (kb)


C

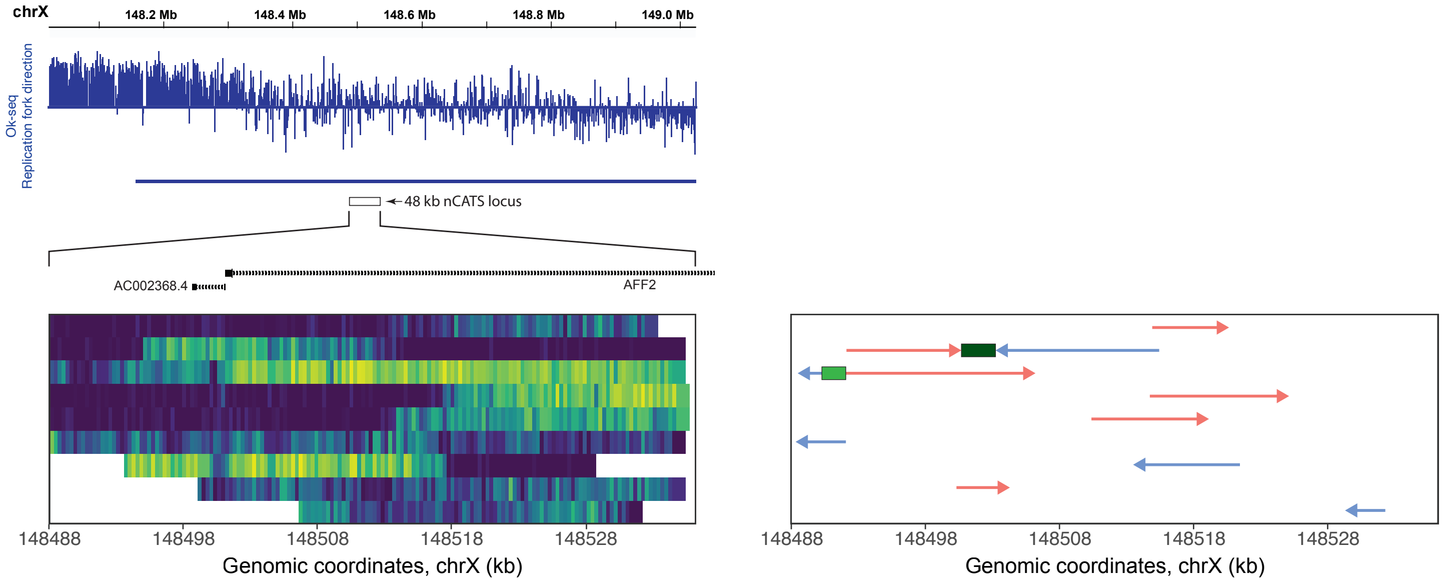

chrX
148.2 Mb
148.4 Mb
148.6 Mb
148.8 Mb
149.0 Mb
Ok-seq
Replication fork direction
48 kb nCATS locus
AC002368.4
AFF2
148488
148498
148508
148518
148528
Genomic coordinates, chrX (kb)
148488
148498
148508
148518
148528
Genomic coordinates, chrX (kb)

### Supplementary Figure S5

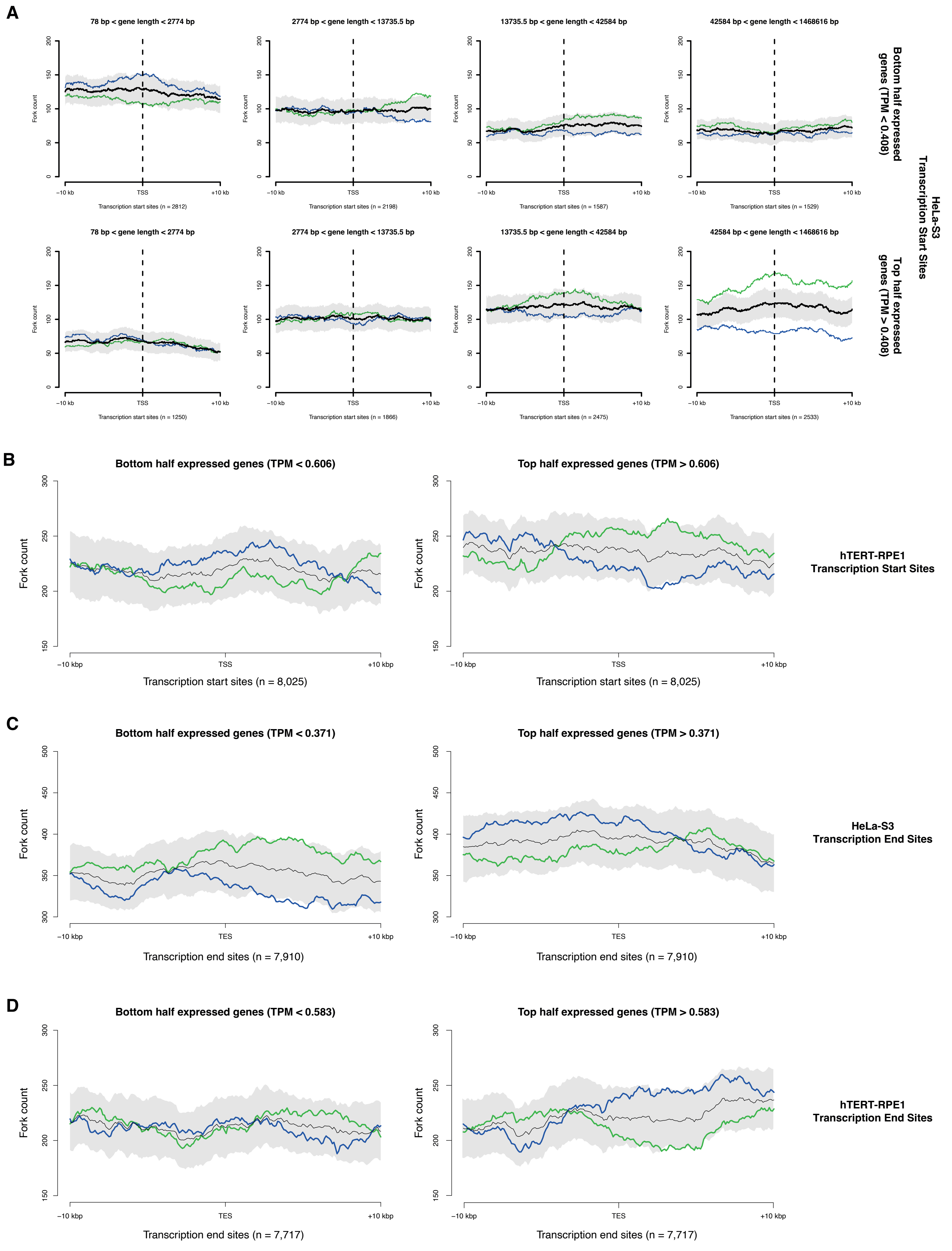

### Supplementary Figure S5

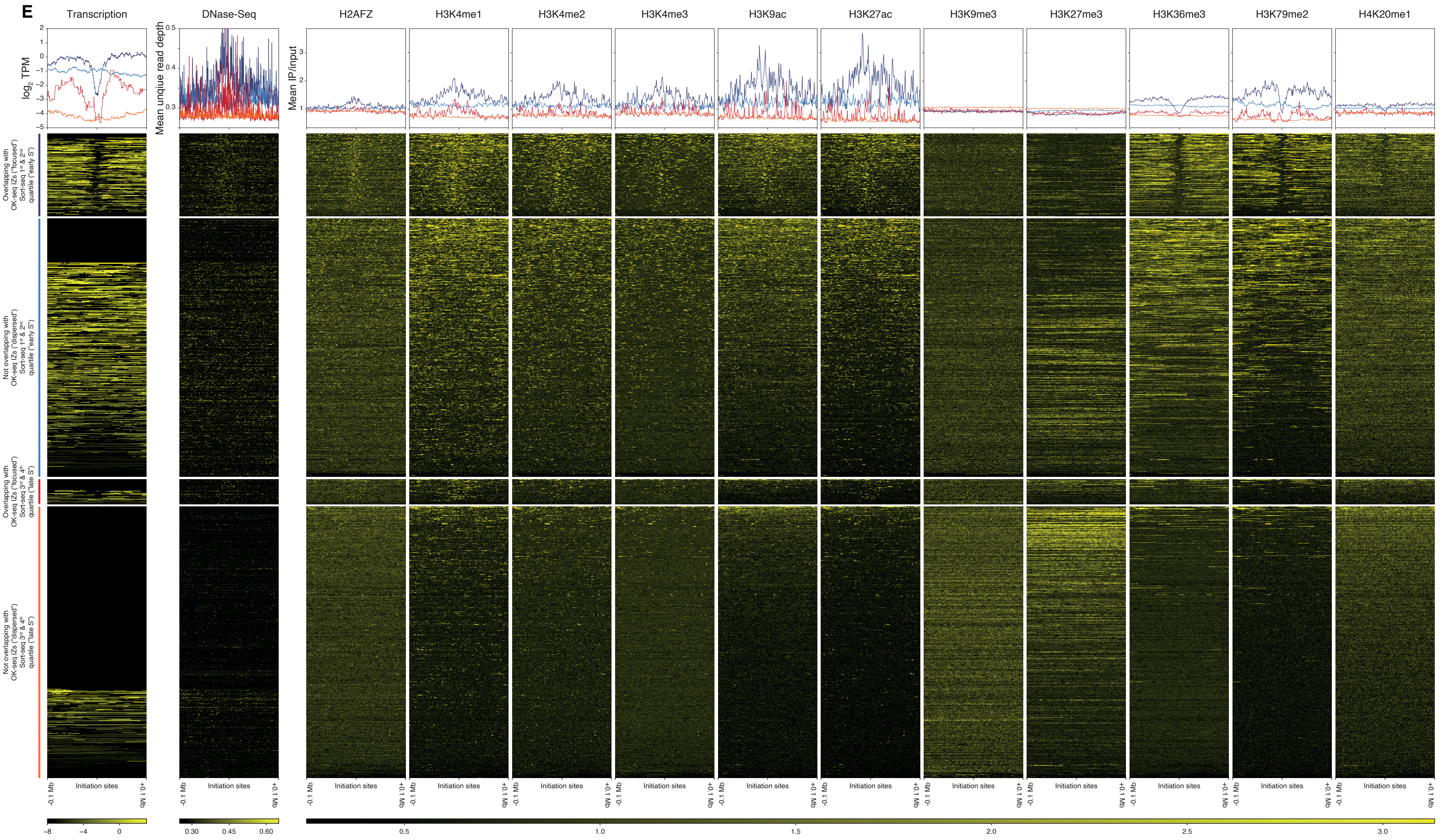
